# A benchmark dataset for individual tree crown delineation in co-registered airborne RGB, LiDAR and hyperspectral imagery from the National Ecological Observation Network

**DOI:** 10.1101/2020.11.16.385088

**Authors:** Ben. G. Weinstein, Sarah J. Graves, Sergio Marconi, Aditya Singh, Alina Zare, Dylan Stewart, Stephanie A. Bohlman, Ethan P. White

## Abstract

Broad scale remote sensing promises to build forest inventories at unprecedented scales. A crucial step in this process is designing individual tree segmentation algorithms to associate pixels into delineated tree crowns. While dozens of tree delineation algorithms have been proposed, their performance is typically not compared based on standard data or evaluation metrics, making it difficult to understand which algorithms perform best under what circumstances. There is a need for an open evaluation benchmark to minimize differences in reported results due to data quality, forest type and evaluation metrics, and to support evaluation of algorithms across a broad range of forest types. Combining RGB, LiDAR and hyperspectral sensor data from the National Ecological Observatory Network’s Airborne Observation Platform with multiple types of evaluation data, we created a novel benchmark dataset to assess individual tree delineation methods. This benchmark dataset includes an R package to standardize evaluation metrics and simplify comparisons between methods. The benchmark dataset contains over 6,000 image-annotated crowns, 424 field-annotated crowns, and 3,777 overstory stem points from a wide range of forest types. In addition, we include over 10,000 training crowns for optional use. We discuss the different evaluation sources and assess the accuracy of the image-annotated crowns by comparing annotations among multiple annotators as well as to overlapping field-annotated crowns. We provide an example submission and score for an open-source baseline for future methods.

## Introduction

Quantifying individual trees is a central task for ecology and management of forested landscapes. Crown delineation is critical for remote sensing of individual trees, as well as improving broad scale studies of forest ecology, silviculture and ecosystem services [1]. There have been dozens of proposed crown delineation algorithms, but these algorithms are designed for and evaluated using a range of different data inputs [2–4], sensor resolutions [5], forest structures [6,7], and evaluation protocols [8,9]. This diversity of approaches makes it difficult to track algorithmic progress and prevents practitioners from weighing tradeoffs in proposed pipelines. For example, [5] recently proposed a pixel-based algorithm for 50 cm pan-sharpened satellite RGB data from a tropical forest in Brazil evaluated against field-collected tree stems. How to compare this to the vector-based algorithm in [10] that uses 10 cm RGB data from fixed-winged aircraft evaluated against image-annotated crowns from oak forests in California? The challenge is compounded by tradeoffs among sensor types. For example, [11] described a graph cut algorithm for LiDAR derived 3D point clouds. Commercial LiDAR data collection can be much more expensive to acquire compared to RGB images, but may yield better delineations. However, it is not yet possible to assess whether any potential improvement in delineation is worth the costs and processing expertise.

One solution to these challenges is to develop a benchmark dataset that can be used to evaluate a wide variety of algorithms and data types. Benchmark datasets are a core tool for knowledge building in data science and have played a vital role in the growth in machine learning [12,13]. A good benchmark dataset has three components: 1) well-curated and open-source data, 2) reasonable evaluation criteria, 3) reproducible and transparent scoring. Authors can then apply their algorithms to the benchmark to create a standardized comparison among methods.

A benchmark dataset for tree crown delineation should balance three objectives. First, co-registered data from multiple sensors is important in evaluating the relative performance of different methods and assisting land managers in planning forest surveys. Second, methods should be able to capture variation in tree forms, either within local forests, or across broad geographic areas. Third, evaluation metrics should have strict training and tests splits that benchmark dataset of individual tree crowns derived from multi-sensor imagery in the National Ecological Observation Network.

### Remote sensing data

The National Ecological Observation Network (NEON) is a large initiative to coordinate data collection across the United States at over 40 geographic sites. Annual data collection includes surveys by the airborne observation platform (AOP) using RGB, LiDAR and hyperspectral sensors (http://data.neonscience.org/), as well as standardized forestry surveys at fixed plots throughout each site. The NEON Airborne Observatory Platform uses fixed-wing aircraft to survey sites during leaf-on-conditions from May-October. Aircraft are flown around 1000m above ground. Sensor data chosen for this benchmark were collected during flights from 2018 and 2019.

### Orthorectified Camera Mosaic (‘RGB’ NEON ID: DP3.30010.001)

The RGB data were acquired with a D8900 camera with a format of 8,984 × 6,732 pixels. Individual images were color rectified, orthorectified and mosaiced to create a single raster image with a pixel size of 0.1 m^2. Mosaic tiles are provided as 1km^2 geoTIFF files and are named based on the utm coordinate at the northwest origin. RGB data have high spatial resolution and individual trees are often visible based on the crown boundary, as well as color differences among individuals due to taxonomy and health status (Figure 1). For more details on NEON camera orthomosaic products see NEON technical document NEON.DOC.005052.

**Figure 1.**
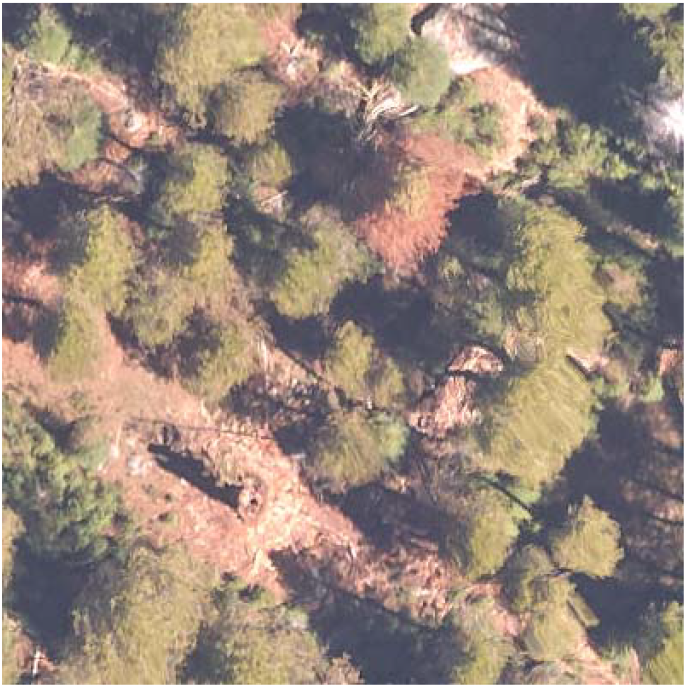
A 40m × 40m evaluation plot of RGB data from the Teakettle Canyon (TEAK) NEON site.

One challenge in creating a multi-sensor dataset is the joint georectification of data types. To ensure spatial overlap between the LiDAR and RGB data, NEON staff overlaid the 0.1m RGB tile on a 1m LiDAR derived surface height model. The difference in spatial resolution can cause some distortion in rectified RGB images. These artifacts are most pronounced at the image edge and were minimized by selecting the centermost portion of each image when creating the RGB mosaic. Some distortion remains and can cause a swirling effect as the image pixels are stretched to match the corresponding LiDAR raster cell. For more information see NEON technical document NEON.DOC.001211vA. We did not include images with large enough distortions to interfere with tree crown detection but kept images with minor distortions to represent the kind of challenging conditions present in applied settings.

### Classified LiDAR Point Cloud (NEON ID: DP1.30003.001)

The LiDAR data are 3D coordinates (4-6 points/m2) that provide high resolution information about tree crown shape and height. LiDAR data are stored as 1km^2^ .laz files (Figure 2). These files contain the x,y,z coordinates for each return, as well as metadata on return intensity and point classification. The LiDAR data has been normalized with respect to classified ground points to standardize height measurements. Tree crowns are often apparent due to gaps among neighboring trees or differences in height among overlapping crowns. For more information on NEON LiDAR data processing see NEON technical document NEON.DOC.001292.

**Figure 2.**
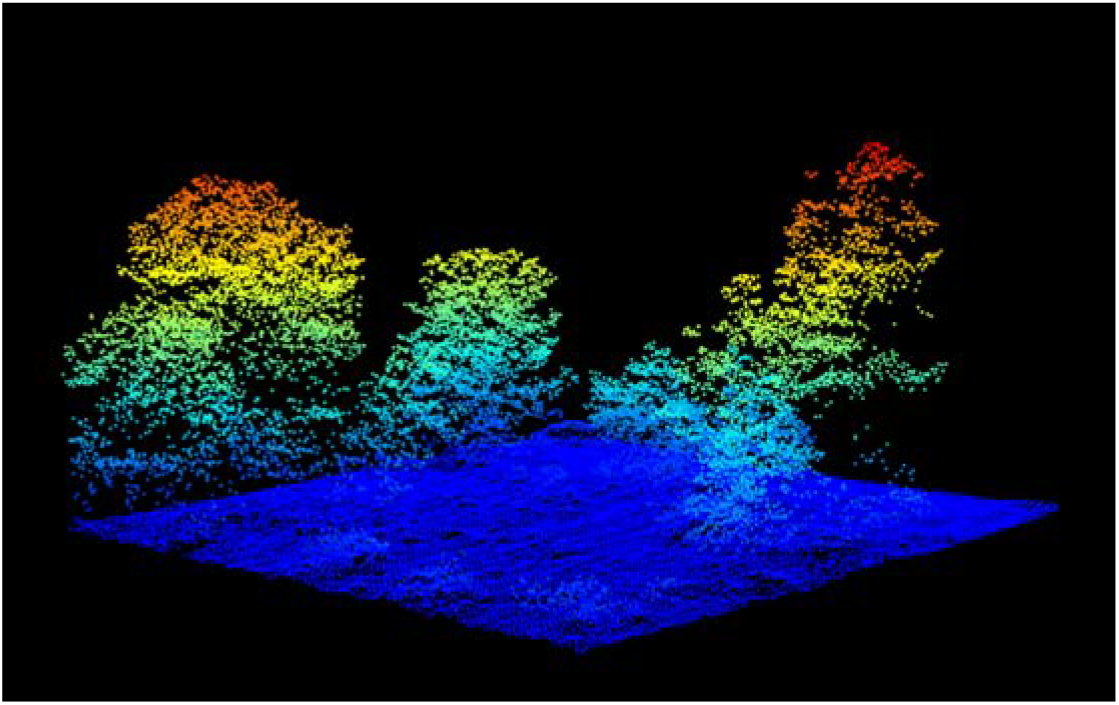
Normalized LIDAR point cloud for evaluation plot SJER_064 from the San Joaquin Experimental Range, California. Points are colored by height above ground.

### Hyperspectral surface reflectance (NEON ID:DP1.30006.001)

NEON’s hyperspectral sensor collects visible and infrared spectrum between approximately 420-2500 nm with a spectral sampling interval of 5nm for a total of 426 bands. NEON provides the orthorectified images with a pixel size of 1 m^2^ in 1 km^2^ tiles that align with the RGB and LiDAR file naming convention. Hyperspectral data, especially in the infrared spectrum, is often used for differentiating tree species (e.g. [14]). In forests with high species diversity, these data may be used to delineate crown boundaries among neighboring trees (Figure 3). All hyperspectral data were collected simultaneously as the RGB data, with the exception of the UNDE site, in which the 2019 RGB data was not available at the time of publication. The 2017 flight was used instead. For more information on hyperspectral data processing and calibration see NEON technical document NEON.DOC.001288.

**Figure 3.**
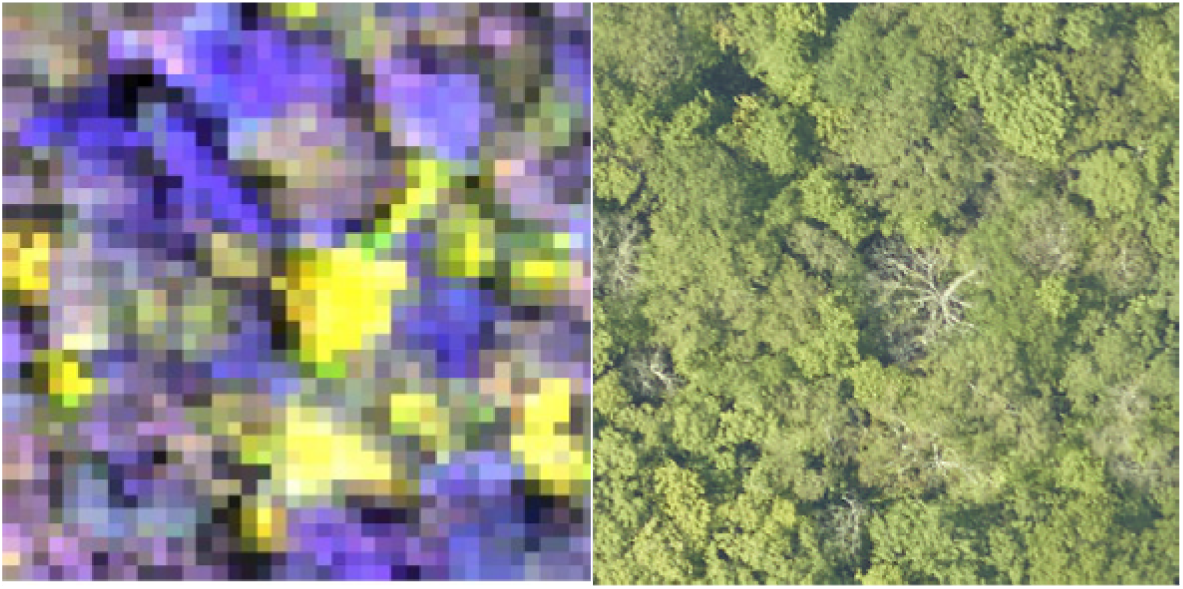
Composite hyperspectral image (left) and corresponding RGB image (right) for the MLBS site. The composite image contains near infrared (940nm), red (650nm), and blue (430nm) channels. Trees that are difficult to segment in RGB imagery (right) may be more separable in hyperspectral imagery due to the differing foliar chemical properties of co-occurring trees.

### Ecosystem Structure (NEON ID: DP3.30015.001)

NEON ‘Ecosystem Structure’ data product is a LiDAR derived height raster at 1m spatial resolution. Often known as a ‘canopy height model’ (CHM), the raster values are the normalized height above ground for each grid cell. For more information on normalization and interpolation to create the raster product see NEON technical document NEON.DOC.002387vA. This data is useful for differentiating crowns in three dimensions, as well as eliminating crowns that are under the 3m threshold used in this benchmark for minimum tree height.

### Woody Plant Vegetation Structure (NEON ID: DP1.10098.001)

Along with sensor data, NEON collects information on individual trees at fixed plots at each NEON site. There are two types of plots included in this dataset: ‘distributed’ plots, which are 20m x 20m fully sampled plots and ‘Tower’ plots, which are 40m × 40m plots with two sampled 20x m 20m quadrants. All trees in sampled areas with a stem diameter of > 10cm are mapped and recorded. For the purposes of this benchmark dataset, the key tree metadata are the stem position, size, and estimated tree height (m).

## Evaluation Data

This benchmark dataset contains the three types of evaluation data: 1) Field-collected stems from 14 sites from the NEON Woody Vegetation Structure dataset, 2) Image-annotated crowns for 22 sites in the NEON network, and 3) Fxield-annotated crowns for two sites in the NEON network. We believe the inclusion of multiple evaluation types is critical because each type of evaluation data has strengths and limitations.

The majority of crown delineation papers assess proposed methods using field-collected stem data in which a single GPS point, representing the position of the basal stem, is used to indicate the position of each tree [15]. There is high confidence that this type of evaluation data represents individual trees, because tree stems are often easy to identify in the field. However, the position of these stems can fail to accurately represent the position of the crown as viewed from above due to a combination of spatial errors in alignment with the image data and the fact that trees can grow at acute angles, such that the center of crowns and position of stems are offset by several meters (Graves et al. 2018). Another limitation of this type of evaluation data is that field-collected stems are typically collected for only a portion of the trees in the landscape. This makes it easy to assess model accuracy (recall), but difficult to assess model precision, since it is not possible to differentiate a non-matching prediction from a correct prediction on a tree without evaluation data. This gap can be critical since an algorithm covering an image in many predictions is likely to have high accuracy, but would be unusable for many scientific applications.

The second type of crown evaluation data are hand-annotated bounding boxes made by an observer looking only at the sensor data [10,16]. We refer to this type of data as ‘image-annotated crowns’. Because this data is annotated by looking directly at the sensor data, there is reduced uncertainty regarding spatial alignment and crown position. It is also relatively easy to scale, allowing data to be collected that cover a wide range of forest types. Since every visible crown in the image can be annotated, both recall and precision can be calculated. However, because this type of evaluation data lacks the presence of a researcher in the field, there can be errors related to the relationship between what appears to be a single crown in the imagery and actual trees. In some cases, what appears to be a single crown in the remote sensing may be composed of multiple smaller trees, or large distinct branches from a single tree may be mistaken for multiple trees.

The final type of evaluation data are ‘field-annotated crowns’ in which an observer annotates the remote-sensing image on a tablet while standing in the field [17]. This approach combines the confidence of the field verification of the field-collected stems with the direct connection to the remote sensing of the image-annotated crowns. The main limitation to this approach is that it is time and labor intensive, which means that only a relatively small amount of validation data can be collected. It is therefore difficult to obtain a large number of crowns across broad enough scales to support evaluation on different forest types. In addition, because of the time investment in collecting these data it is rare for these data to annotate all trees in a region, there are similar issues to field-collected stem data for assessing model precision.

### Evaluation Methods

#### Field-collected stems

The field-collected stems were filtered from the raw NEON Woody Vegetation Structure data to reflect that crowns were likely to be visible in the canopy. Understory tree detection is an important avenue of future work, but is beyond the scope of this benchmark. For more information on algorithmically selecting overstory trees see S1. Using the overstory stem data, the goal is to predict a single bounding box that matches to a single field-validated stem. For each field plot we score the proportion of field stems that fall within a single predicted crown. Field stems can only be assigned to one prediction, such that if two predictions overlap a single field stem, only one is considered a positive match. The resulting proportion of stems with a positive match can be used to estimate the stem recall rate, ranging from 0 (no correctly matched stems) to 1 (all stems are matched).

### Image-annotated Crowns

We selected airborne imagery from 22 sites surveyed by the NEON airborne platform. The evaluation sites were chosen based on the availability of the three types of sensor data, as well as to represent a breadth of forest landscapes across the continental US. For each site we annotated a minimum of two 40mx40m evaluation plots. The location of these evaluation plots mirrored the sampling design of the NEON Woody Vegetation structure ‘tower’ plots.

Images were annotated using the program RectLabel (Table 1). For each visible tree, we created a bounding box (xmin, ymin, xmax, ymax) that covered the tree crown (Figure 4). The coordinates of this box are relative to the top left corner of each image.

**Table 1.** Benchmark evaluation scores for the DeepForest python package. The submission data are provided with the NeonTreeEvaluation package installation. The recall and precision range from 0 to 1. Recall is the proportion of true positives divided by the total number of samples. Precision is the proportion of predictions that are true positives.

| Image-annotated |  | Field-annotated | Field collected |
| --- | --- | --- | --- |
| Crowns |  | Crowns | Stems |
| Recall | Precision | Recall | Recall |
| 79.0 | 65.9 | 72.2 | 74.0 |

**Figure 4.**
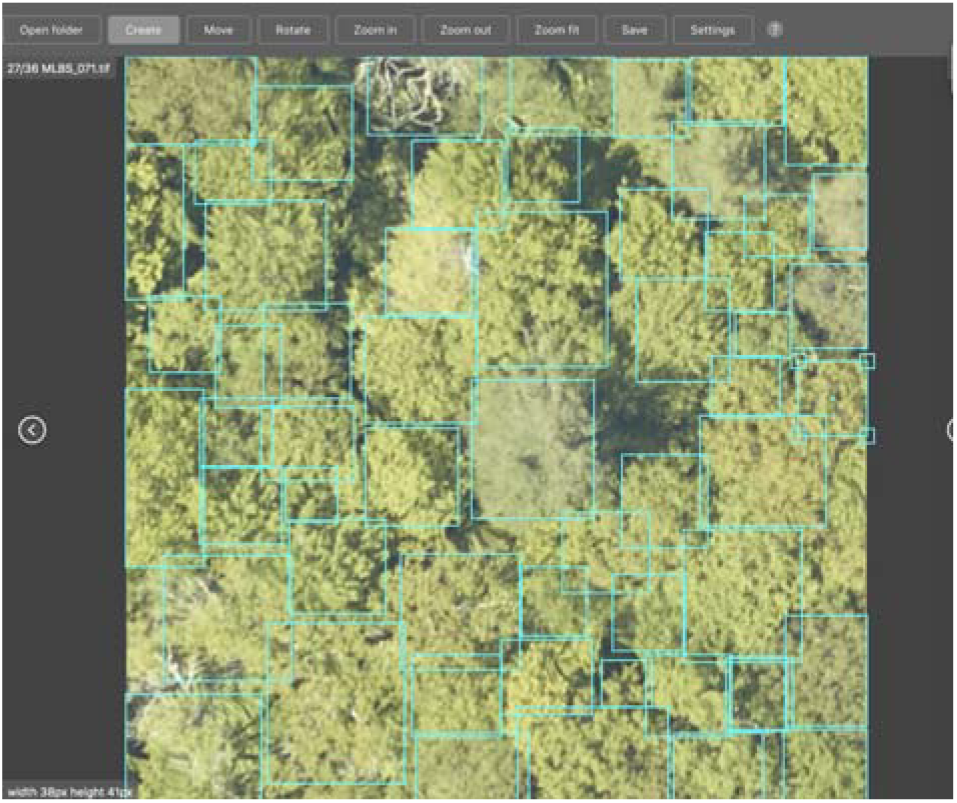
Screenshot of the program RectLabel used for tree annotation for the image-annotated crowns for NEON plot MLBS_071. For each visible tree crown, we created a four point bounding box.

We carefully annotated the evaluation images by comparing the RGB, LiDAR and hyperspectral data. Using all three products made it possible to more accurately distinguish neighboring trees in images by simultaneously assessing visual patterns (RGB), using variation in spectral signatures to distinguish different species (hyperspectral), and looking at the three dimensional structure of the tree (LiDAR). For some sites, such as OSBS, the crowns were most visible in the LiDAR height model, whereas for closed canopy sites such as MLBS, the hyperspectral and RGB data were most useful. When working with the hyperspectral data we primarily used a composite three band hyperspectral image containing near infrared (940nm), red (650nm), and blue (430nm) channels, which showed good differences between neighboring trees of different types (Figure 5d, h). We also augmented the RGB data to view subtle changes in pixel values using a decorrelation stretch (Figure 5b, f). To enforce a minimum size threshold for tree annotations, we removed any vegetation less than 3m in height. Annotations were also compared to the woody vegetation structure field data wherever possible. Each evaluation plot overlaps with a NEON 40m × 40m plot. Within each of these plots, NEON field crews survey a 20×20 subplot, and therefore while field data are available for most plots in the dataset, they do not cover every tree in the image. The woody vegetation structure data contains information on field estimated height and maximum crown diameter for the majority of field collected stems. We annotated all trees, regardless of health status, provided they were standing at the time of data collection. To facilitate labeling at broad extents, tree crowns in this dataset are simplified into rectangular bounding boxes rather than convex polygons. This is in line with the majority of object detection tasks in computer vision, but can be limiting when delineating fuzzy crown boundaries.

**Figure 5.**
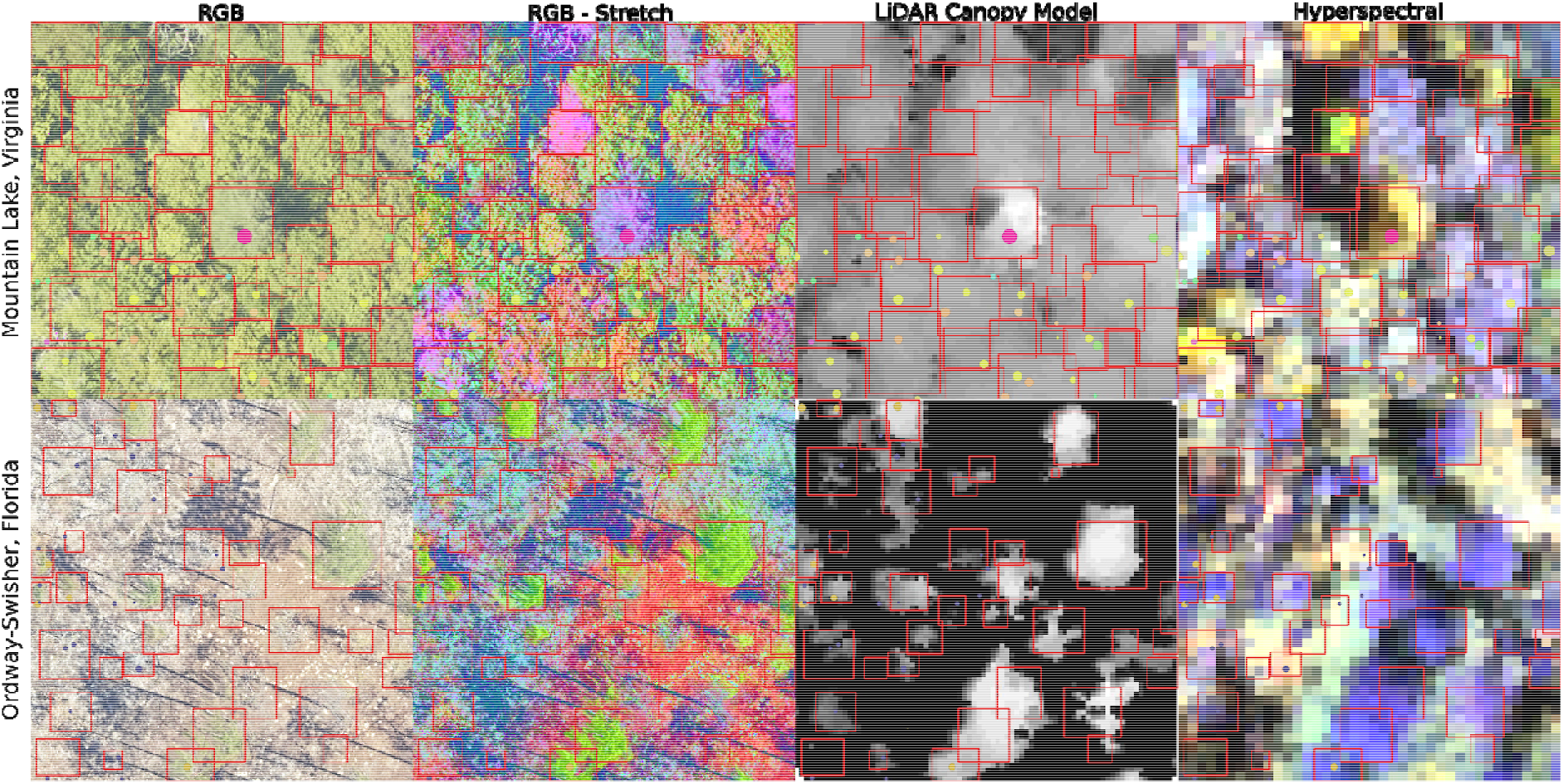
Tree crown annotations for the evaluation data set for two sites in the National Ecological Observation Network. Using the RGB, LiDAR and hyperspectral products together contributes to more careful crown annotation. For some sites, such as MLBS (top row), the RGB and hyperspectral data are useful for differentiating overlapping crowns. For other sites, such as OSBS (bottom row) the LiDAR point cloud, shown as a rasterized height image, is most useful in capturing crown extent. The RGB-stretch image was produced by transforming the RGB data in the three principal components space. To create a three-band hyperspectral image, we used channels from the red, blue and infrared spectrum to capture changes in reflectance not apparent in the RGB imagery.

To convert overlap among predicted and ground truth bounding boxes into measures of accuracy and precision, the most common approach is to compare the overlap using the intersection-over-union metric (IoU)(e.g. Lin et al. 2017). IoU is the ratio between the area of the overlap and the area of the combined bounding box region. The metric ranges between 0 (no overlap) to 1 (perfect overlap). We considered boxes which have an IoU score of greater than 0.4 as true positive, and scores less than 0.4 as false negatives (Figure 6). A single threshold was selected to be able to get an overall summary performance statistic. The 0.4 value was chosen based on visual evaluation of the threshold that indicated a good visual match between the predicted and observed crown (see ‘uncertainty in annotations’). In the wider computer vision literature, the conventional threshold value for overlap is 0.5 (e.g. [20]), but this value is arbitrary and does not ultimately relate to any particular ecological question. We tested a range of overlap thresholds from 0.3 (less overlap among matching crowns) to 0.6 (more overlap among matching crowns) and found that 0.4 balanced a rigorous cutoff without spuriously removing trees that would be useful for downstream analysis.

**Figure 6.**
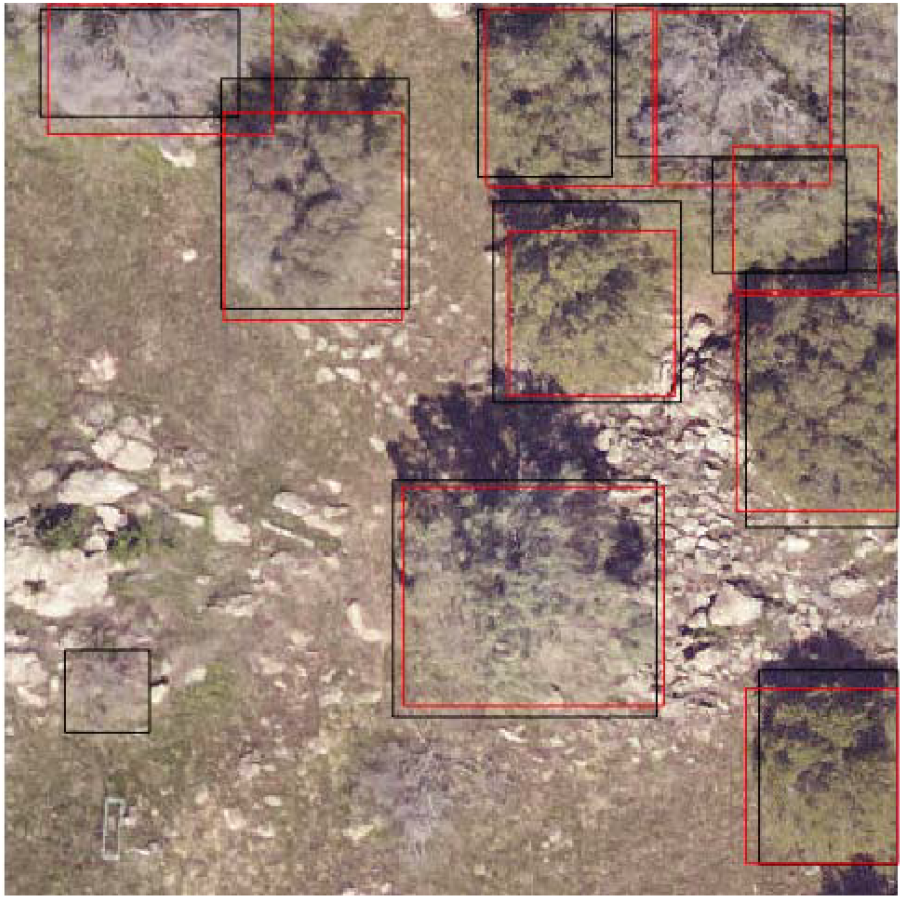
Example evaluation from the NeonTreeEvaluation R package. Predicted boxes (see below) in red and ground truth boxes are in black. In this image there are 10 image-annotated boxes, and 9 predictions. Each prediction matches an image-annotated box with an intersection-over-union score of greater than 0.4. This leads to a recall score of 0.9 and a precision score of 1.

Using this overlap threshold we calculated recall, defined as the proportion of bounding boxes correctly predicted, and precision, defined as the proportion of predictions that matched a ground truth bounding box. If multiple predictions overlapped a single ground truth box, we matched the prediction of highest overlap to the ground truth box. Predictions that did not overlap with any ground truth are considered false positives. To create a single summary statistic, we used the mean precision and recall per image rather than pooling results across sites. We chose this statistic to emphasize the wide geographic variance in tree conditions. Pooling results together would be biased towards performance in forests with higher tree density at the expense of forests with lower tree density.

### Field-annotated Crowns

Individual trees were mapped by visiting two NEON sites and directly mapping polygonal tree crowns in the remote sensing images using a field tablet and GIS software while looking at each tree from the ground [18]. False-color composites from the hyperspectral data, RGB, and LiDAR canopy height images were loaded onto tablet computers that were equipped with GPS receivers. While in the field, researchers digitized crown boundaries based on the location, size, and shape of the crown seen in the field. Trees were mapped in 2014 and 2015, and all polygons were manually checked against the most recent NEON imagery. All crowns that were no longer apparent in the RGB or LiDAR data (due to tree fall, or overgrowth) were removed from the dataset, and minor adjustments to crown shape and position were refined after examining multiple years of RGB imagery. No adjustments to the polygons were made due to crown expansion.

Field-collected crowns are spaced widely throughout the forest with only a handful overlapping in a single 40mx40m image. Therefore only recall (the proportion of predictions that match a field-annotated crown polygon at an intersection-over-union threshold of greater than 0.4) can be assessed. Precision cannot be assessed, since it is not possible to differentiate an incorrect prediction from a tree that was not mapped.

### Training Annotations

It is common for computer vision benchmarks to include fixed training and testing data. During our research on crown delineation algorithms (Weinstein et al. (2019, 2020a, 2020b) we annotated many geographic tiles separate from the evaluation data. The training sites were selected to capture a range of forest conditions including oak woodland (NEON site: SJER), mixed pine (TEAK), alpine forest (NIWO), riparian woodlands (LENO), southern pinelands (OSBS), and eastern deciduous forest (MLBS). The training tiles were chosen at random from the NEON data portal, with the requirement that they did not contain a large amount of missing data (e.g. due to the presence of an edge of a site) and they did not overlap with any evaluation plots. Depending on the tree density at the site, we optionally cropped the 1 km to a smaller size to create more tractable sizes for annotation. This data is released as part of the benchmark dataset. However, our goal is to promote the best possible crown-delineation algorithm regardless of training data, and so we do not believe the inclusion of this training data should preclude others from applying their trained algorithms on the evaluation dataset. Given the large size of training tiles, they were less thoroughly reviewed and were only checked in the RGB imagery.

## Uncertainty in annotations

### Differences between image-only annotators

Since the image-annotated crowns were done by visually inspecting the images, the exact position and number of bounding boxes in an image will depend on the annotators’ interpretation of crown location. Image interpretation is a standard practice for creating validation sets in remote sensing (e.g. [19]), but depends on the skill of the interpreter and always introduces uncertainty to validation. In many computer vision tasks, class boundaries are clear and definitive. However, the combination of image quality, spatially overlapping crowns, as well as the two-dimensional view of a complex three-dimensional canopy, makes it difficult to always identify where one crown ends and another begins. To assess this uncertainty, a 2nd observer annotated 71 evaluation plots using the same data as the primary annotator. We then compared these annotations using a range of intersection-over-union (IoU) thresholds to indicate true positive matching crowns (Figure 7). We found that crown recall among annotators ranged from approximately 70% at lower IoU thresholds to 90% at higher IoU thresholds. This variance indicates that differences between annotators reflect differences in crown extent, not differences in whether or not a tree is present. If tree detection was the primary area of disagreement, changing the IoU threshold would have minimal effect on the recall and precision rates. This was also supported at the plot level, where the number of trees and mean tree height from the LiDAR cloud were nearly identical across multiple annotators, but there was more variation in the mean crown area (Figure 7).

**Figure 7.**
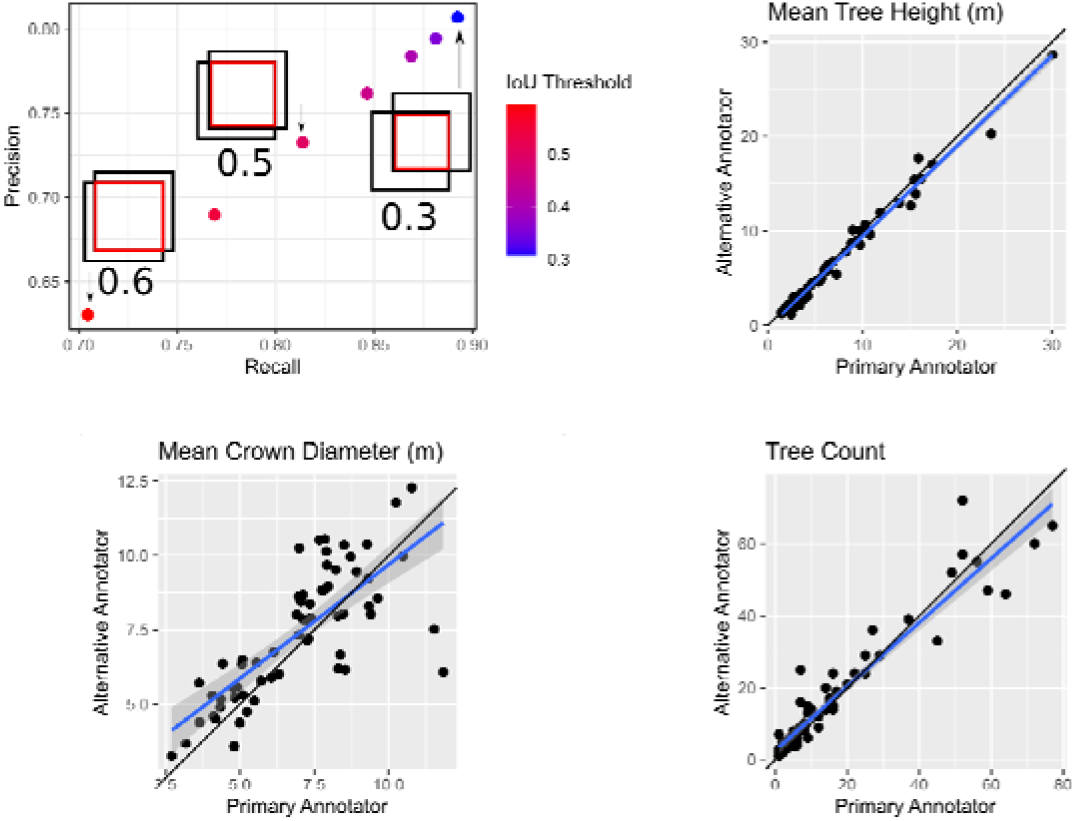
Intersection-over-union scores (top left), as well as plot in level inferences, between the primary annotator and a 2nd annotator. For the IoU scores, we plotted precision and recall for 7 different intersection-over-union thresholds. As the overlap threshold decreases, the two annotators tend to agree on ground truth tree crowns. Analysis is based on 71 evaluation images (n=1172 trees) that were separately annotated by two different annotators.

### Comparison among image-annotated and field-annotated crowns

To assess the ability for image-annotated crowns to represent field validated data, we compared image-annotation made by the primary annotator (BW) with the field-annotated crowns (annotated by SG) at two sites for which there was overlapping remote sensing imagery. We compared image annotations and field crowns using the crown recall rate, defined as the proportion of field-annotated crowns that overlap a image-annotated crowns (IoU threshold > 0.4), and the stem recall rate, defined as the proportion of field-annotated crown centroids that are within a single image-annotated bounding box. The primary annotator independently annotated 1553 crowns in images that overlapped with 91 field collected crowns at Mountain Lake Biological Station (MLBS) and 27 crowns at Ordway-Swisher Biological Station (OSBS). To prevent the annotator identifying the obvious location of the field crown, the test image encompassed a large area. Using field-annotated crowns as ground truth, the image annotations had a centroid rate of 96.7% indicating that image annotation can identify the presence of trees in all but rare cases. There was more disagreement in the extent of crown boundaries. The image-annotated crowns had a crown overlap recall of 78.0% with the field-annotated crown polygons. Visual inspection of more stringent intersection-over-union thresholds showed a disconnect between the quantitative score and the qualitative assessment of the image-annotated performance (Figure 8), supporting the use of a 0.4 threshold for identifying a match with empirical tree crowns. However, this decision is subjective and will depend on the sensitivity of downstream analysis using predicted crowns. While we anticipated greater recall for large field-annotated crowns, we found only a modest pattern between increased crown area of field-annotated crowns and correct image-annotated match. In general, errors tend to be marginally biased towards oversegmentation, where large crowns are divided into smaller sets sets of branches, but both types of errors occur in relatively similar frequencies (Figure 9).

**Figure 8.**
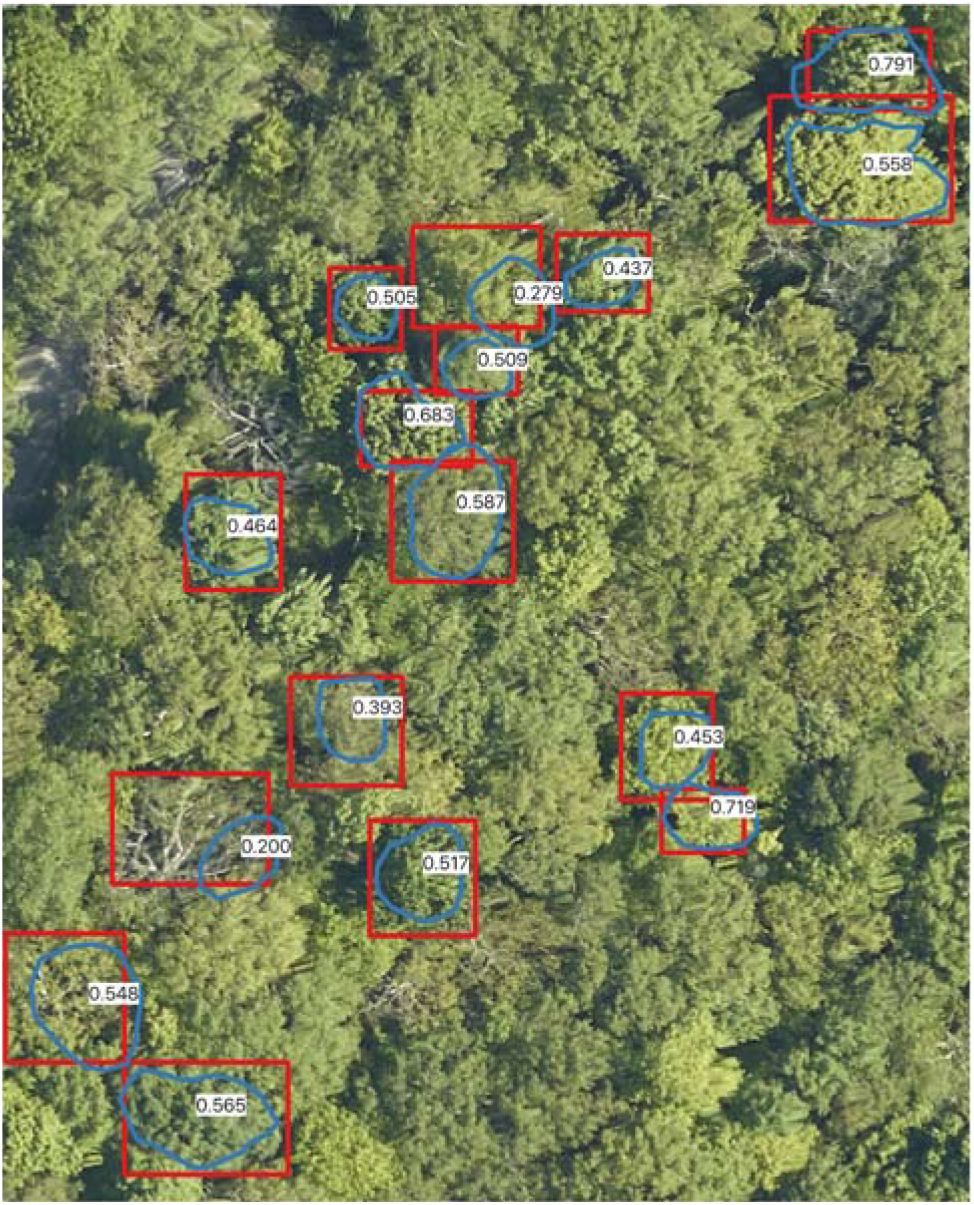
Comparison of field-annotated crowns made by one author (SG) in blue (n=16) and image-annotated crowns made by another author (BW) in red at Mountain Lake Biological Station, Virginia. Intersection-over-union scores are shown in white. Only the image-annotated crowns associated with the field crowns are shown (out of the 206 image-annotated crowns in this image). From this and similar visualizations we determined that a threshold of 0.4 was a reasonable choice for eliminating crowns that are not sufficiently overlapping to be used for ecological analysis.

**Figure 9.**
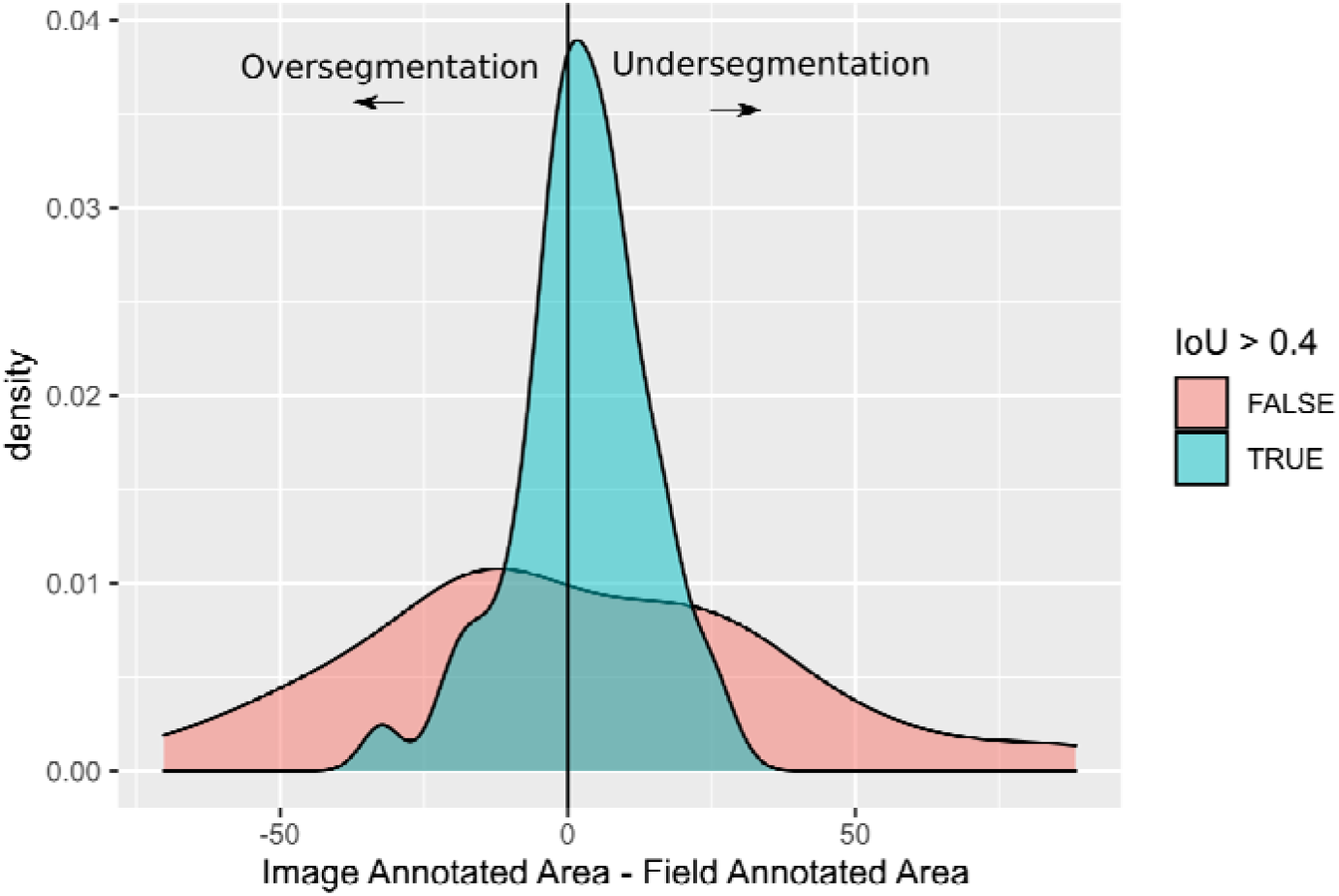
Comparison of the crown bounding box area of matched field-annotated and image-annotated crowns. Oversegmented crowns have image-annotations that are too small for the field-annotated crowns. This typically occurs when large trees are erroneously split into smaller crowns. Undersegmented crowns have image-annotations that are too large for the field-annotated crowns. This typically occurs when dense stands of trees are combined into a single crown. On average the image-annotations that are below the true positive intersection-over-union threshold of 0.4 (in red) tend to be oversegmented. The bounding box area of the field-annotated crowns was used instead of polygons to reduce the difference in annotation format and focus on oversegmentation versus undersegmentation in crown detection.

### NeonTreeEvaluation R Package

To maximize the value of the benchmark dataset and standardize evaluation procedures, we developed an R package (https://github.com/weecology/NeonTreeEvaluation_package) for downloading the evaluation data and running the evaluation workflows. This package takes a standard submission format as input and can be extended to include additional evaluation statistics. This reproducible workflow will be key in creating a more transparent process for future comparisons among tree crown detection algorithms. This repo also contains a leaderboard for users to submit scores and submission documents for future reproducible analysis.

To demonstrate the performance of the benchmark, we used the recently published DeepForest python package to predict crowns in the evaluation data [21]. DeepForest is a RGB deep learning model that predicts tree crown bounding boxes. The prebuilt model in DeepForest was trained with the training data described above, but did not use or overlap spatially with any evaluation data in the benchmark. For more information on the deep learning approach see [10,16,21]. Following the best practices for computational biology benchmarking described in [13], we emphasize that the DeepForest algorithm was designed in conjunction with these evaluation data and it is therefore not surprising that it performs well, with image-annotated boxes and field-annotated crown polygons both at approximately 70% accuracy (Table 1, Figure 10). It is also notable that despite the uncertainty with the crown area of the image-annotated crowns, the overall score is similar among data types.

**Figure 10.**
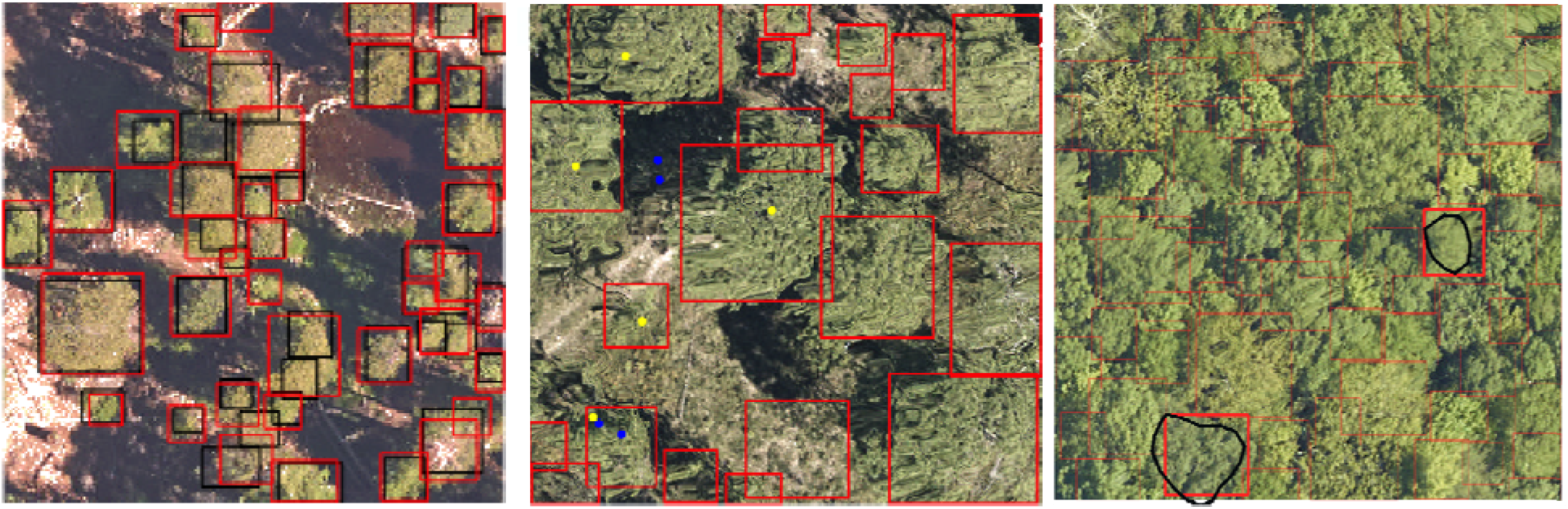
Example predictions using the DeepForest algorithm. A) DeepForest predictions in red and compared to image-annotated crowns in black from Teakettle Canyon, California. B) DeepForest predictions in red are compared to field-collected stems, with matching stems in yellow and missing stems in blue, from Jones Ecological Research Center, Georgia. C) DeepForest predictions in red with the field-annotated crown in black from Mountain Lake Biological Station, Virginia. The matching prediction is shown in bold while the other predictions are faded for visibility.

## Discussion

The benchmark data set is designed to assess tree crown detection in airborne imagery of forests. This dataset differs from standard computer vision benchmarks due to the large number and high variability of objects in the dataset. There are over a hundred trees in some images, which vary in size, shape and spectral properties. Generalization across geography is a fundamental challenge in remote sensing, and this dataset is constructed to study potential tradeoffs between local accuracy and generalizability to unseen forest conditions. Even when algorithms have been developed solely for one site, generalization can be important for future users in adapting them to new forest conditions.

This dataset is the first to include aligned data from RGB, LiDAR and hyperspectral sensors for a range of geographic areas. While they are most often analyzed separately, each of these data types may be useful for tree crown detection. Three-dimensional LiDAR data has high spatial resolution, but it can be difficult to identify tree boundaries due to a lack of spectral information. RGB data has spectral information but lacks context on vertical shape and height. Hyperspectral data is useful for differentiating tree species but is generally at a coarser spatial resolution. Combining sensor data may lead to more robust and generalizable models of tree detection at broad scales, which makes having all three data types aligned an important component of a forward-looking benchmark dataset.

While the annotations are represented by 2D bounding boxes, there is significant opportunity to extend the benchmark dataset into new formats and dimensions. For example, there has been recent interest in object detection using input rasters, both as a replacement for traditional bounding boxes, and as an additional step in refining pixel-based contours of object boundaries. By rasterizing the annotated bounding boxes, the dataset can be used to compare segmentation strategies such as raster-based versus regional proposal networks. Furthermore, combining 2D optical data and 3D point cloud annotations remains an active area of model development. Trees have complex 3D and 2D representations and the data provided in this benchmark could be used to develop new evaluation procedures across dimensions.

By providing a repeatable evaluation workflow, we hope to reduce the uncertainty in novel algorithm development and promote model and data sharing among researchers. Initial work in [16] showed that deep learning algorithms can learn from multiple geographies simultaneously, without losing accuracy on the local forest type. This means that data sharing among researchers can provide mutual benefit to all applications, even from disparate forest types. By standardizing evaluation criteria, we hope to foster collaboration and comparative studies to improve the accuracy, generalization, and transparency of tree crown delineation.

## Acknowledgements

We would like to thank NEON staff and in particular Tristan Goulden and Courtney Meier for their assistance and support. This research was supported by the Gordon and Betty Moore Foundation’s Data-Driven Discovery Initiative (GBMF4563) to E.P. White and by the National Science Foundation (1926542) to E.P. White, S.A. Bohlman, A. Zare, D.Z. Wang, and A. Singh. The funders had no role in study design, data collection and analysis, decision to publish, or preparation of the manuscript.

## S1. Selecting overstory trees for the NEON field-collected stems NEON

The following filters were applied to the raw NEON field data (ID) after download. An overstory reference tree must have

- Valid spatial coordinates
- A unique height measurement per sampling period. Species double recorded but with different heights were discarded
- Sampled in more than one year to verify height measurement
- Changes in between year field heights of less than 6m
- Classified as alive
- A minimum height of 3m to match the threshold in the remote sensing workflow.
- Be at least within 5m of the canopy as measured by the LiDAR height model extracted at the stem location. The was used to prevent matching with understory trees in the event that overstory trees were eliminated due to failing in one of the above conditions, or not sampled by NEON.

To match trees we took the closest height when two predictions and field stems overlapped. dropped CLBJ since only 3 points met this criteria. All other sites did not have any data that met this criteria.

## S2. List of annotations for each geographic site

**Table 1.**
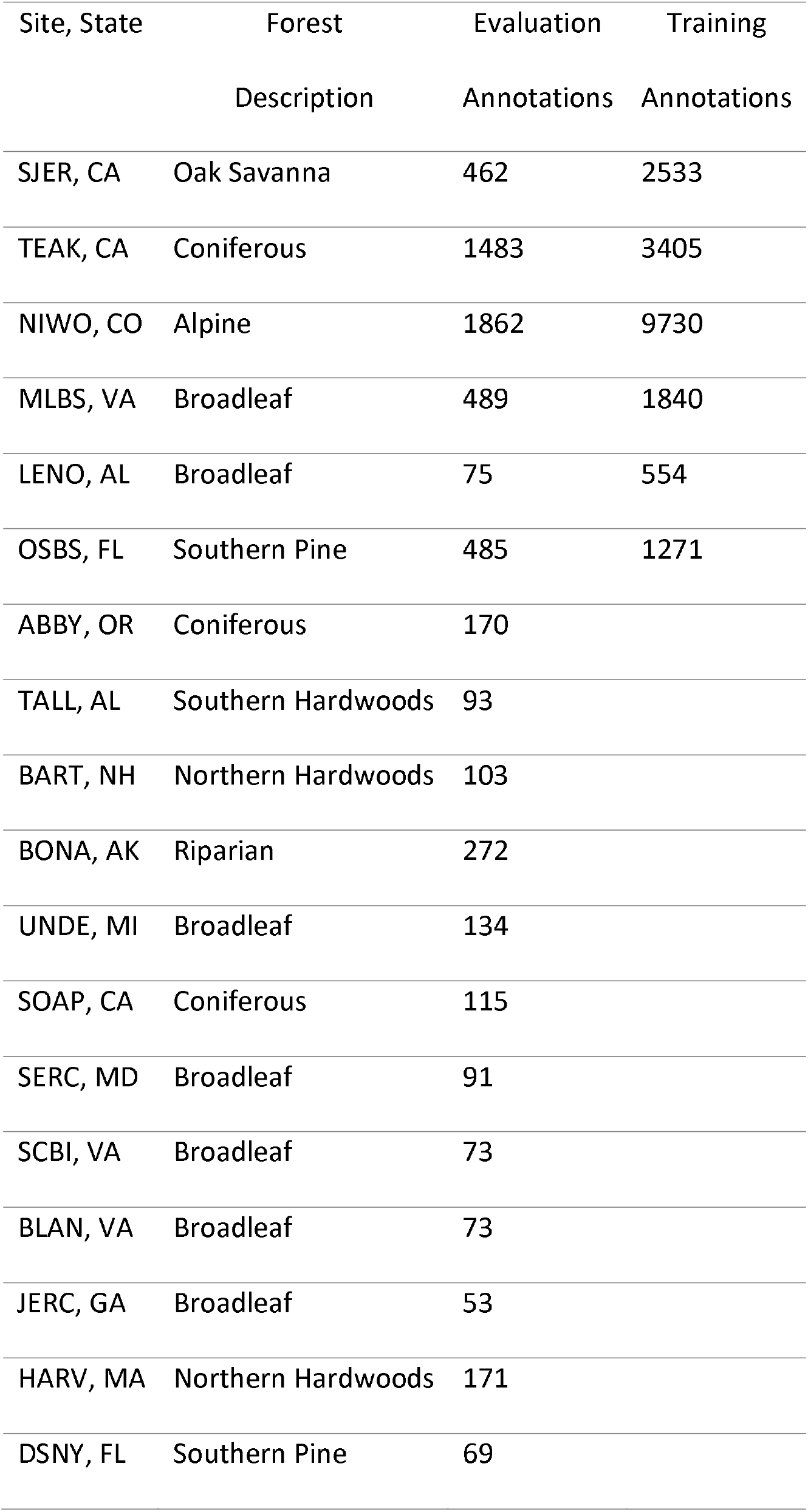
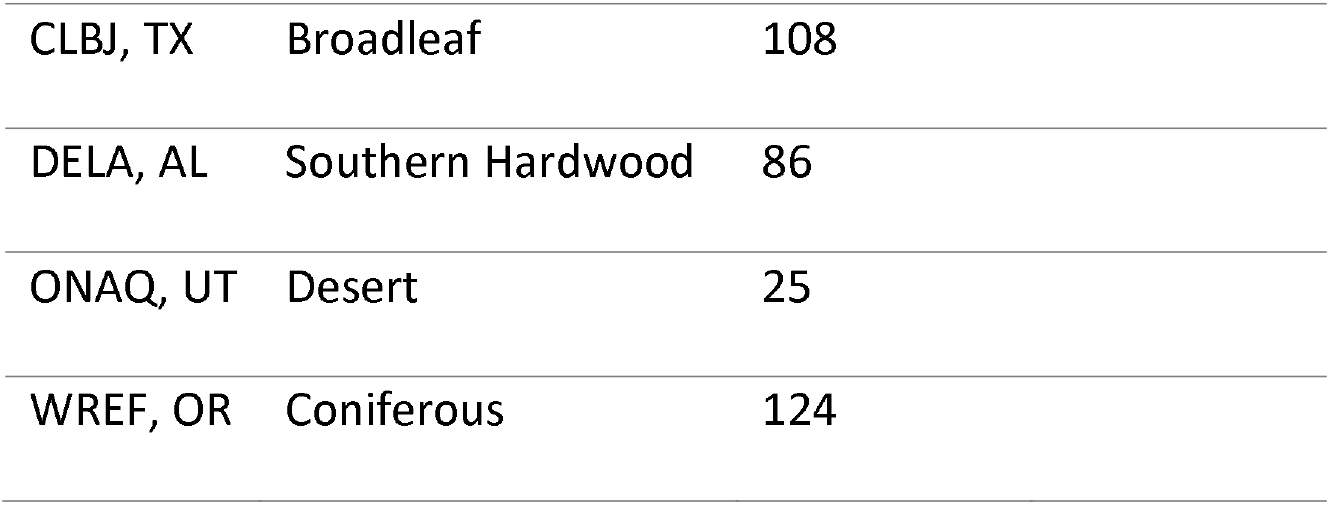
The number of image-annotated tree crowns for each site.

| Site, State | Forest | Evaluation | Training |
| --- | --- | --- | --- |
|  | Description | Annotations | Annotations |
| SJER, CA | Oak Savanna | 462 | 2533 |
| TEAK, CA | Coniferous | 1483 | 3405 |
| NIWO, CO | Alpine | 1862 | 9730 |
| MLBS, VA | Broadleaf | 489 | 1840 |
| LENO, AL | Broadleaf | 75 | 554 |
| OSBS, FL | Southern Pine | 485 | 1271 |
| ABBY, OR | Coniferous | 170 |  |
| TALL, AL | Southern Hardwoods | 93 |  |
| BART, NH | Northern Hardwoods | 103 |  |
| BONA, AK | Riparian | 272 |  |
| UNDE, MI | Broadleaf | 134 |  |
| SOAP, CA | Coniferous | 115 |  |
| SERC, MD | Broadleaf | 91 |  |
| SCBI, VA | Broadleaf | 73 |  |
| BLAN, VA | Broadleaf | 73 |  |
| JERC, GA | Broadleaf | 53 |  |
| HARV, MA | Northern Hardwoods | 171 |  |
| DSNY, FL | Southern Pine | 69 |  |

|  |  |  |
| --- | --- | --- |
| CLBJ, TX | Broadleaf | 108 |
| DELA, AL | Southern Hardwood | 86 |
| ONAQ, UT | Desert | 25 |
| WREF, OR | Coniferous | 124 |

